# RNA Polymerase II Degradation Triggered by DNA Repair Occurs *In Trans* and Independently of how the Lesion is Recognised

**DOI:** 10.1101/2024.08.30.610509

**Authors:** Ramveer Choudhary, Juan Cristobal Muñoz, Inés Beckerman, Giulia Bastianello, León Alberto Bouvier, Marco Foiani, Manuel J. Muñoz

## Abstract

In response to DNA damage, RPB1, the catalytic subunit of RNA Polymerase II (RNAPII), is degraded by the ubiquitin-proteasome system. Degradation models only consider transcriptionally engaged molecules, where a stalled RNAPII complex functions as a lesion recognition factor and its RPB1 subunit is proposed to be subsequently degraded to facilitate access of core Nucleotide Excision Repair (NER) factors. This Transcription Coupled repair is complemented by the Global Genome repair (GG-NER) system, where lesions are recognized by the XPE and XPC factors. Here we show that RPB1 degradation is controlled *in trans* by a pathway that depends on NER activity, irrespectively of whether the lesion is recognized by RNAPII itself or by GG-NER factors. Incomplete lesion repair due to absence of any core NER factor enhances RPB1 degradation, indicating that the signal controlling RPB1 abundance is started by lesion recognition and continues until DNA repair is completed. Consistent with an *in trans* mechanism, damage-induced RPB1 degradation is not restricted to active nor phosphorylated RPB1 molecules and depends on Cullin-RING ubiquitin ligases. These findings uncover a repair-dependent mechanism controlling RPB1 levels and provide a rationale for the control of gene expression under stress, where more damage implies more repair and less RPB1 levels, hence restricting RNAPII activity.

## Introduction

Gene expression and lesion repair, two of the major DNA-based molecular mechanisms in a cell, necessarily interact with each other. This idea is well illustrated by the Transcription Coupled-Nucleotide Excision Repair system (TC-NER), where lesion-stalled RNA polymerases initiate DNA repair (Nieto Moreno et al., 2023; van den Heuvel et al., 2021). At the time of acute DNA damage induction, as when exposing cells to ultraviolet (UV) light, every active RNAPII complex not only transcribes a nascent RNA, but also serves as a DNA lesion scanner. As a direct consequence, repair of template strands in transcriptionally active genes is faster than in the rest of the genome (Bohr et al., 1985; Hu et al., 2017). Conversely, lesions in coding strands and transcriptionally silent regions are recognized by the xeroderma pigmentosum (XP) factors XPC and XPE, also known as DDB2, which act as lesion recognition factors in the other branch of NER, the Global Genome-NER (GG-NER) (Apelt et al., 2021). After lesion recognition by either RNAPII stalling or XPC-XPE, both pathways converge to favor the recruitment of core NER factors involved in the actual repair. These factors include XPB and XPD helicases, part of the general transcription factor TFIIH, that together with XPA scan the DNA to verify the presence of a lesion. In such a case, recruitment of the endonucleases XPF and XPG to excise the damaged strand is favored, and the single-stranded DNA (ssDNA) gap is filled by DNA synthesis and ligation (Cohen & Adar, 2023; Spivak, 2015).

RPB1, the major and catalytic subunit of RNAPII, is not only unique because of its repetitive Carboxy Terminal Domain (CTD), which is subject to multiple regulatory post-translational modifications, but also because it is specifically degraded in response to different DNA damaging agents. As an example, exposure to UV light induces the proteasomal degradation of RPB1 (Ratner et al., 1998), as well as deep changes in gene expression (M. J. Muñoz et al., 2009, 2017; Williamson et al., 2017). Connecting these findings, identification of the lysine residue involved in RPB1 degradation revealed that RNAPII levels shape the gene expression response, since cells with impaired RPB1 ubiquitylation and degradation showed altered gene expression patterns and affected viability in response to UV light (Nakazawa et al., 2020; Tufegdžić Vidaković et al., 2020).

Although it is clear that RNAPII stalling is a mean for lesion detection and also that RPB1 is specifically degraded in response to DNA damage, the connection between these two, damage detection and RPB1 degradation, is not obvious. The last resort model suggests that the stalled RNAPII molecule that acts as a lesion recognition factor is the one that is degraded (Wilson et al., 2013). *In situ* degradation of lesion stalled RPB1 would be needed to clear the lesion while simultaneously favoring the recruitment of core NER factors (XPA, XPB, XPD, XPF and XPG) to induce repair. Nevertheless, there is no evidence showing that the actual RBP1 molecule stalled in front of a lesion is the one that is degraded by the proteasome, nor which are the pathways or mechanisms involved in the control of RPB1 levels.

Beyond the last resort model and the idea of *in situ* degradation, recent evidence suggested that promoter-proximal paused RPB1 molecules can be degraded in vicinity of the lesion, but not at the lesion (Bay et al., 2022; Steurer et al., 2022). Although with different proximity to damaged DNA, these models propose degradation of phosphorylated and transcriptionally engaged RNAPII molecules, implying a direct link between an active RNAPII complex and RPB1 degradation. Nonetheless, since transcription is inhibited after damage (Rockx et al., 2000), the idea of a transcription-dependent mechanism controlling the prominent decrease in RPB1 levels is difficult to reconcile.

Apart from RPB1 degradation and TC-NER mediated repair, DNA lesions also induce GG-NER. In fact, a possible connection between GG-NER and RPB1 degradation has not been addressed, being currently unknown if these two events are coupled or concurrent, i.e., just happening at the same time. Furthermore, since lesion recognition, not only by GG-NER but also by TC-NER, precedes repair, it is plausible that total NER activity leads to the degradation of any given RPB1 molecule.

Using human keratinocytes exposed to UV light and other DNA damaging agents, here we show that total NER-mediated repair controls RBP1 degradation and, therefore, RNAPII activity. Considering that TC-NER is at its peak immediately after damage, while GG-NER is active for longer periods of time (Hu et al., 2017), RPB1 degradation is controlled in a sequential manner, first by TC-NER and then by GG-NER. Furthermore, incomplete lesion repair due to the absence of any core NER factor (XPA, XPB, XPD, XPF and XPG) enhanced RPB1 degradation, suggesting that signaling for RPB1 degradation is initiated by lesion recognition, either by TC-NER or GG-NER, and continues until DNA repair is completed. In agreement with an *in trans* signaling pathway controlling RPB1 levels, we present evidence suggesting that in the context of TC-NER, one RPB1 molecule, part of an active RNAPII complex, signals another RPB1 molecule for degradation. Finally, we show that transcriptionally inactive RPB1 molecules can be targeted for degradation in a Cullin-RING E3 ligase dependent manner, indicating that a NER-dependent pathway regulates, *in trans*, RPB1 abundance.

## Results

Given the central role of RNAPII in gene expression, it is expected that RPB1 abundance is of outmost importance for proper cell function. In this sense, datasets regarding variations in gene copy number and human disease prevalence have been recently published (Collins et al., 2022). To analyze the impact of RPB1 levels in human disease, we used these datasets to study the effect of increased or decreased dosage of *POLR2A*, the gene encoding RPB1. As expected, results in Figure S1A showed that alterations in the amount of *POLR2A* are highly associated with the onset of pathologies. Nevertheless, and despite its relevance, mechanisms controlling RPB1 abundance are largely unknown.

### TC-NER controls RPB1 levels in human keratinocytes

Prominent variations in RPB1 levels have been historically observed under an acute response to DNA damage, where RPB1 abundance decreases due to specific proteasome-dependent degradation. To better understand the mechanisms controlling RPB1 levels and having in mind that current degradation models rely on phosphorylated and transcriptionally engaged RNAPII molecules, we decided to force lesion recognition by active RNAPII complexes by inactivating GG-NER. To this end, we generated an XPC and XPE double knock out (dKO) human keratinocyte cell line (Figure S1B), deficient in damage recognition and repair by GG-NER (Figure S1C). Keratinocytes, the most common cell type in human skin, are naturally exposed to UV light and can be easily arrested in G1 by serum withdrawal (Figure S1D). To avoid excessive cell death and activation of pathways due to conflicts between DNA and RNA polymerases in S-phase, we based our experimental set up in G1 arrested human keratinocytes (HaCaT cells) (Boukamp et al., 1988) exposed to UV light. As expected, UV exposure of G1 arrested wildtype (WT) HaCaT cells induced the proteasome dependent degradation of RPB1 (Figure S1E).

To confirm the widespread idea that TC-NER can induce RPB1 degradation, and also that GG-NER deficient cells are TC-NER proficient (van Hoffen et al., 1995), we analyzed RPB1 levels in total extracts of both, WT and XPC-XPE dKO GG-NER deficient HaCaT cells. As expected, results in Figure 1A showed a comparable reduction in RPB1 levels shortly after UV irradiation (∼3 h) of WT and dKO cells. To confirm these results, we treated WT and dKO cells with Illudin S, a DNA alkylating agent that, unlike UV-induced Cyclobutane Pyrimidine Dimers (CPDs) and 6-4 Pyrimidine-pyrimidone Photoproducts (6-4PPs), can be recognized exclusively by TC-NER (Jaspers et al., 2002). Results in Figure 1B show that exposure of WT and dKO cells to Illudin S induced a comparable decrease in RPB1 levels in both cell types, thus confirming that TC-NER can induce RPB1 degradation and also that GG-NER deficient cells are TC-NER proficient.

**Figure 1.**
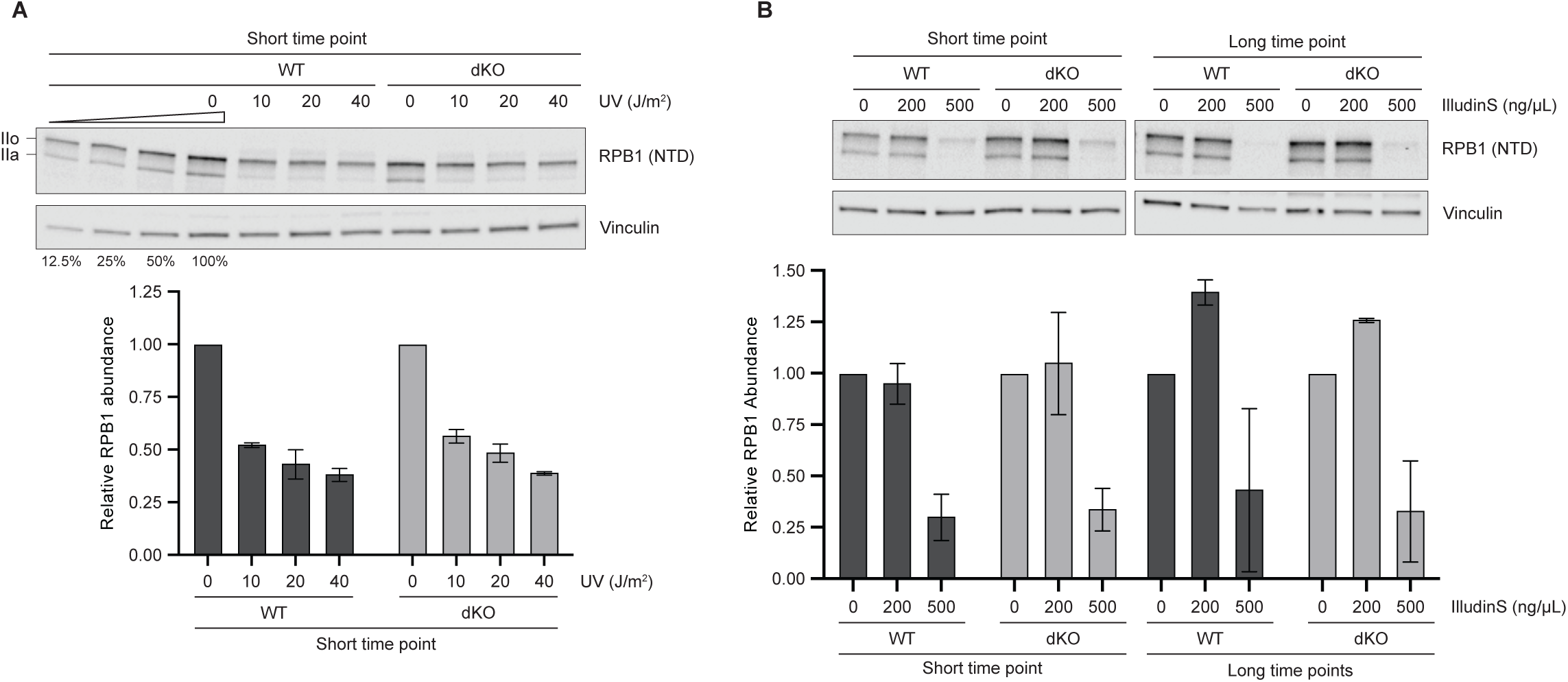
TC-NER controls RPB1 levels in human keratinocytes. (A) Gl arrested WT and XPC-XPE double CRISPR KO (dKO) HaCaT human keratinocytes were irradiated with the indicated doses of UV light and harvested 3 h after (short time point). Relative RPBl abundance was assessed by western blot using antibodies against the N-Terminal Domain (NTD) of RPBl and Vinculin as a loading control. IIo and IIa indicates the phosphorylated and unphosphorylated forms of RPBl. Images of a representative experiment and mean ± SEM of RPBl abundance relative to O J/m^2^ condition from two independent experiments are shown. (B) Gl arrested WT and XPC-XPE double KO (dKO) HaCaT human keratinocytes were treated with the indicated doses of Illudin S and harvested 3 h (short time point) or l2 h (long time point) after. Relative RPBl abundance was assessed as before. Images of a representative experiment and mean ± SEM of RPBl abundance relative to O J/m^2^ condition from two independent experiments are shown.

### GG-NER controls RPB1 levels in human keratinocytes

Given that TC-NER induces not only RPB1 degradation but also DNA repair, we wondered whether total NER, or just TC-NER, could modulate RPB1 levels. To analyze a possible role for NER-mediated repair in the control of RPB1 levels, we recognized that any TC-NER contribution should occur shortly after UV irradiation, where TC-NER activity accounts for most of NER-mediated DNA repair. Moreover, at longer time points after damage induction, when TC-NER activity is barely detectable (Hu et al., 2017), GG-NER deficient cells will lack any NER mediated repair, providing an opportunity to assess a possible role for NER-dependent repair in the control of RPB1 levels. To this end, we measured RPB1 levels in GG-NER proficient (WT) and deficient (dKO) cells ∼12 h after UV irradiation, where TC-NER is no longer active. Results in Figure 2A showed higher levels of RPB1 in GG-NER deficient than in WT cells, pointing towards a role for total NER-mediated DNA repair in the control of RPB1 levels. Comparable results were obtained using single XPC or XPE knockout cells (Figure S2A) as well as siRNA-mediated knockdown of XPC in WT cells (Figures S2B and S2C). These results strongly suggest that NER-mediated repair controls RPB1 abundance. Moreover, since TC-NER activity is maximum immediately after UV exposure, while GG-NER deals with UV-induced DNA damage for much longer, these results favor a model of sequential control of RPB1 levels, first by TC-NER and then GG-NER.

**Figure 2.**
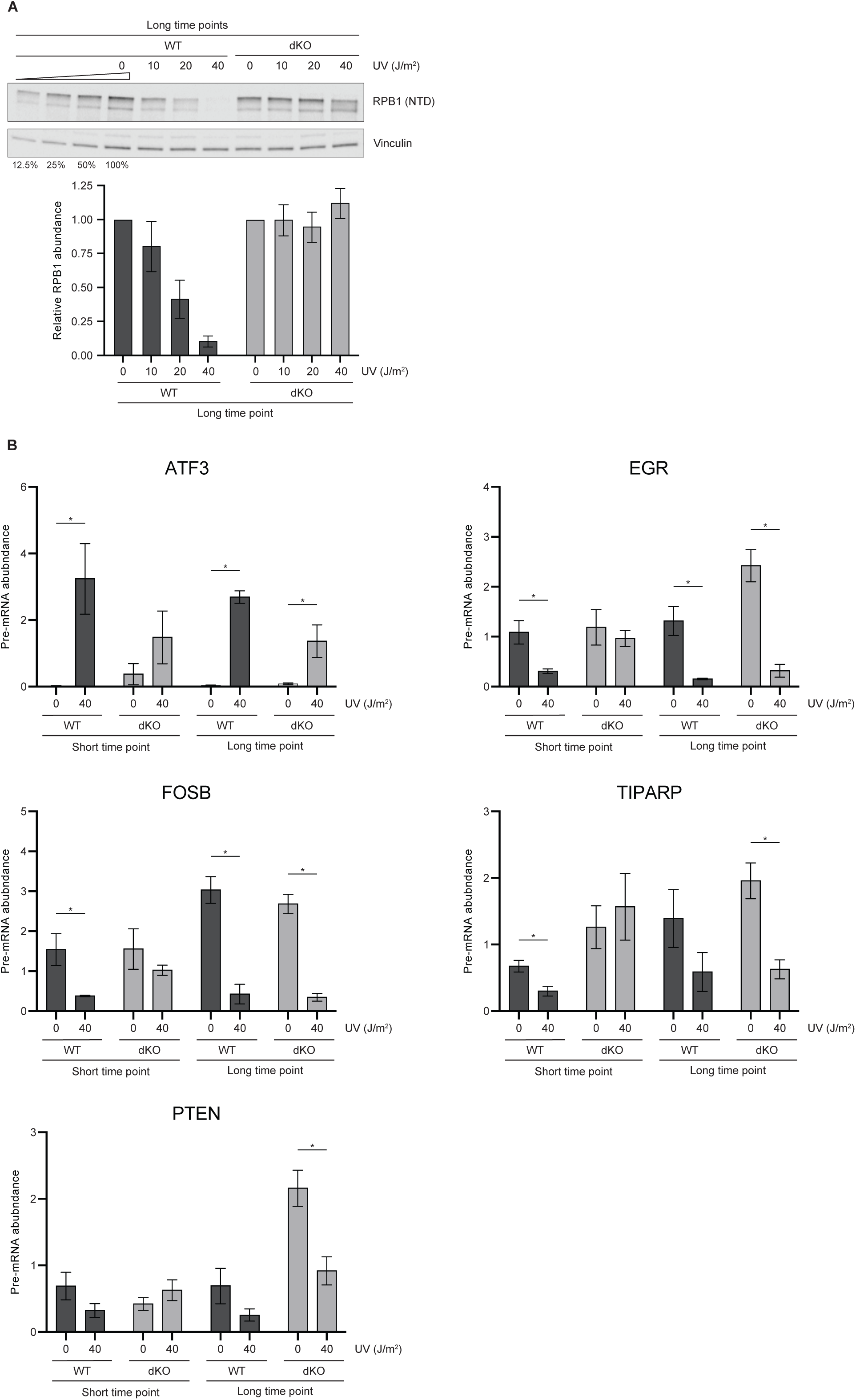
GG-NER controls RPBl levels in human keratinocytes. (A) Gl arrested WT and XPC-XPE double KO (dKO) HaCaT human keratinocytes were irradiated with the indicated doses of UV light and harvested l2 h after (long time point). Relative RPBl abundance was assessed as before. Images of a representative experiment and mean ± SEM of RPBl abundance relative to 0 J/m^2^ condition from two independent experiments are shown. (B) Gl arrested WT and XPC-XPE double KO (dKO) HaCaT cells were treated with the indicated doses of UV light and harvested after 3 h (short time point) or l2 h (long time point) for total RNA preparation. Pre-mRNA levels of the indicated individual gene were assessed by RT-qPCR relative to U6 mRNA levels (housekeeping). Data are shown as mean ± SEM. Statistically significant differences are indicated with an asterisk (p < 0.05, unpaired Student’s t test). n=3-4

Having in mind that RPB1 levels were found to affect the gene expression response to damage (Nakazawa et al., 2020; Tufegdžić Vidaković et al., 2020), and also that GG-NER deficient cells have altered RPB1 levels (Figures 2A, S2A and S2B), we decided to analyze gene expression in WT and GG-NER deficient HaCaT cells. To this end, we treated WT and dKO cells with UV light and measured ongoing transcription by analyzing a set of pre-mRNAs by RT-qPCR. As expected, results in Figure 2B showed that UV-treated GG-NER deficient cells have altered gene expression patterns when compared to WT cells, in agreement with previous reports showing altered gene expression in XPC mutant fibroblasts exposed to UV light (Andrade-Lima et al., 2015). In line with this, total RNA recovery synthesis showed higher levels of overall expression in dKO than in WT cells (Figure S2D). Therefore, we conclude that GG-NER deficient human keratinocytes display increased RPB1 levels and RNAPII activity in response to DNA damage.

### NER-mediated DNA repair controls RPB1 degradation *in trans*

Results in Figure 1 and Figure 2 suggest that the repair process controls, by an unknown mechanism, RPB1 abundance. To deepen into a possible role for NER in the control of RPB1 levels, we next generated an XPA knockout HaCaT cell line (XPA KO) (Figure S3A), in which damaged DNA is recognized but the repair process cannot be completed. In comparison to WT keratinocytes, XPA KO HaCaT cells showed lower levels of RPB1 in response to UV exposure (Figure 3A). The enhanced reduction in RPB1 levels upon UV proved to be proteasome-dependent (Figure S3B) and was also observed in WT cells transfected with siRNAs against XPA (Figure S3C). These results suggest that the NER system activates a pathway that, in turn, favors RPB1 ubiquitylation and its subsequent proteasomal degradation.

**Figure 3.**
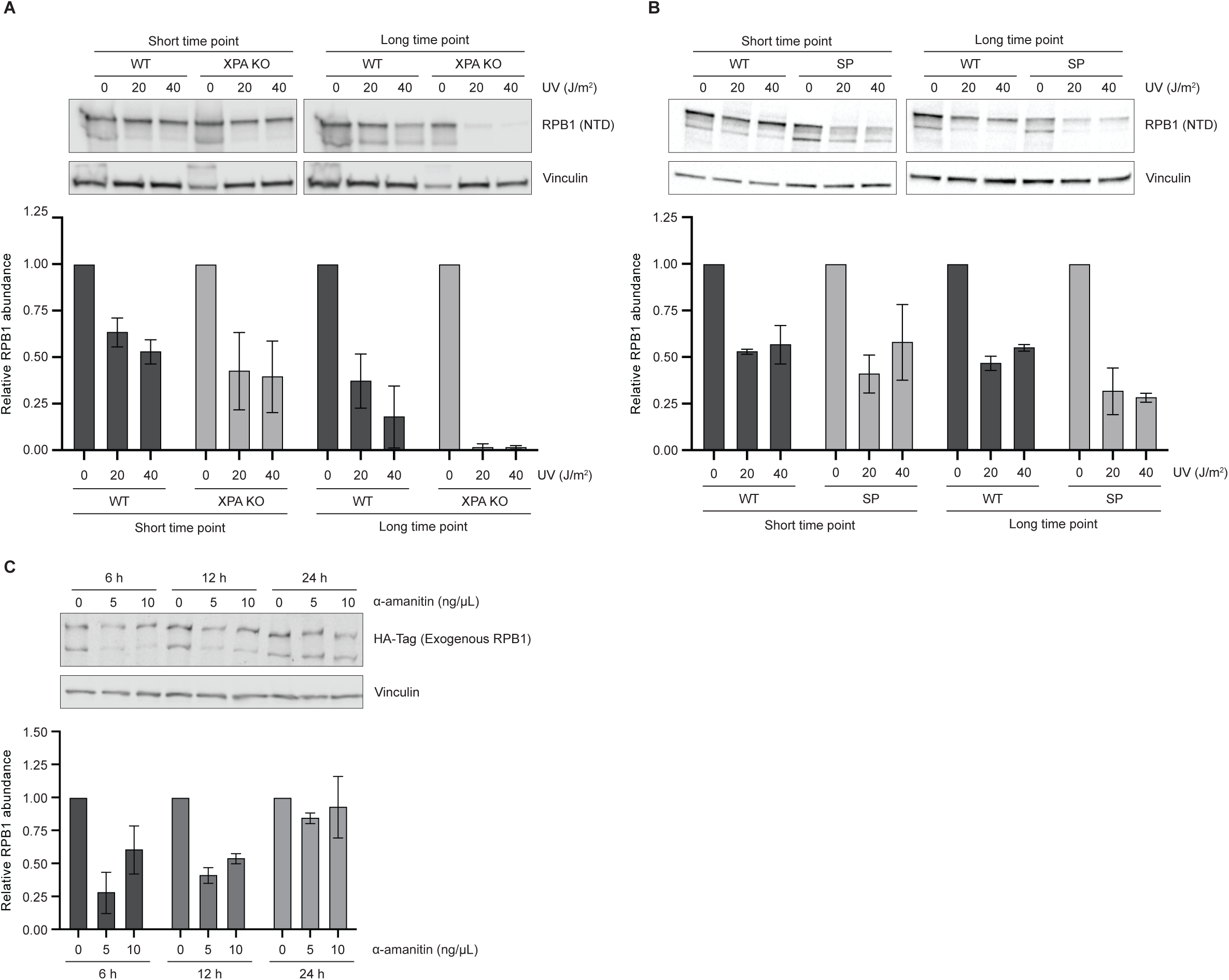
NER-mediated DNA repair controls RPBl degradation in trans. (A) G1 arrested WT and XPA KO HaCaT human keratinocytes were irradiated with the indicated doses of UV light and harvested 3 h (short time point) or 12 h (long time point) after. Relative RPB1 abundance was assessed as before. Images of a representative experiment and mean ± SEM of RPB1 abundance relative to O J/m^2^ condition from two independent experiments are shown. (B) G1 arrested WT HaCaT human keratinocytes were incubated for 1 h with Spironolactone (SP, 2.5 µM) to induce XPB knockdown. Cells were then irradiated with the indicated doses of UV light and harvested 3 h (short time points) or 12 h (long time point) after. Relative RPB1 abundance was assessed as before. Images of a representative experiment and mean ± SEM of RPB1 abundance relative to O J/m^2^ condition from two independent experiments are shown. (C) HEK293T human cells were transfected with a plasmid encoding an HA-tagged and a-amanitin resistant version of RPB1. 48 h after transfection, cells were treated with the indicated doses of a-amanitin and harvested after 6 h, 12 h or 24 h. Relative levels of the HA-tagged and a-amanitin resistant version of RPB1 were assessed by Western bot using antibodies against the HA tag and Vinculin as a loading control. Images of a representative experiment and mean ± SEM of RPB1 abundance relative to O ng/µL of a-amanitin condition from two independent experiments are shown.

To rule out specific functions of XPA, we next treated WT HaCaT cells with Spironolactone (SP), a drug that induces degradation of XPB (Alekseev et al., 2014), another core NER repair factor. In agreement with a role for NER in the control of RPB1 abundance, results in Figures 3B, S3D and S3E showed that treatment with SP enhanced the UV-induced degradation of RPB1. Next, we inhibited the gap-filling step of the NER reaction by treating G1 arrested WT HaCaT cells with the DNA polymerase inhibitor Aphidicolin. Results in Figure S3F showed that after UV exposure, RPB1 levels were further reduced in cells where repair cannot be completed, in this case by a compromised gap-filling step. In line with this, similar results were obtained when transfecting WT HaCaT cells with siRNAs directed against other core NER repair factors, namely XPD, XPF and XPG (Figures S3G and S3H). Altogether, these results suggest that DNA repair by the NER mechanism controls RPB1 levels in human keratinocytes, where less lesion recognition implies less RPB1 degradation (Figure 2) and an incomplete repair process results in enhanced degradation levels (Figure 3). Of note, RPB1 degradation in UV-treated keratinocytes was barely affected by inhibiting classical DNA damage response kinases, such as ATM, ATR, DNA-PK and GSK3 (Figures S3I and S3J).

The fact that there are no RPB1 molecules directly involved in GG-NER suggests the presence of a repair-dependent pathway controlling, *in trans*, the levels of RPB1. Consequently, since TC-NER can induce RPB1 degradation (Figure 1), it is possible that one RPB1 molecule, part of a lesion-stalled RNAPII complex, favors degradation of a distinct RPB1 molecule. To address this matter, we reasoned that the experimental limitation was to be able to distinguish RPB1 molecules involved in either lesion recognition or targeted for degradation. To address this issue, we considered α-amanitin, a drug that impairs ribonucleotide incorporation by sensitive RPB1 molecules inducing RNAPII stalling, RPB1 degradation, and changes in gene expression patterns, similarly to UV irradiation (Bao et al., 2024; Bernecky et al., 2016; Brueckner & Cramer, 2008). Having in mind that α-amanitin resistance can be achieved by a point mutation in RPB1’s coding sequence (N792D), this drug has been widely used to replace the endogenous RPB1 pool with an exogenous and resistant version (Nguyen et al., 1996). Cells transfected with α-amanitin resistant RPB1 expression plasmids have, initially, two populations of RPB1 molecules: α-amanitin sensitive, encoded in the endogenous loci, and α-amanitin resistant, exogenous molecules. After adding α-amanitin to the culture medium, gene expression is typically assessed 24 to 48 h later, when the endogenous RPB1 pool is barely detectable and the exogenous version is evident. In the context of an *in trans* degradation pathway, we hypothesized that shortly after adding α-amanitin, a sensitive and stalled RPB1 molecule might induce, through a TC-NER reaction, degradation of either sensitive or resistant molecules. To test this idea, we transfected HEK293T cells with an α-amanitin resistant RPB1 expression plasmid and then analyzed the levels of both, α-amanitin sensitive and resistant pools in response to α-amanitin treatment. In line with a NER-dependent *in trans* degradation mechanism, results in Figures 3C, S3K and S3L show that the resistant pool is in fact partially and initially sensitive to α-amanitin but later, and in agreement with the many studies that took advantage of α-amanitin to exchange different versions of RPB1, the resistant pool is stabilized while the sensitive pool is further degraded. In this matter, we hypothesize that after adding the drug, the levels of the sensitive pool, able to target sensitive or resistant RPB1 molecules for degradation, will decrease with time. Conversely, as sensitive molecules progressively decrease, the α-amanitin resistant pool stabilizes, reaching a maximum that will ultimately depend on *de novo* expression of sensitive RPB1 molecules encoded in the endogenous loci.

### Transcriptionally inactive RPB1 molecules can be targeted for degradation

Either induced by TC-NER or GG-NER, RPB1 levels are being controlled by a NER-dependent *in trans* pathway. We next wondered about the nature of the targeted RPB1 molecules, part of either transcriptionally active or inactive RNAPII complexes. To this end, we firstly used 5,6-dichloro-1-β-D-ribofuranosylbenzimidazole (DRB), a CTD’s kinase inhibitor that favors the unphosphorylated and transcriptionally inactive isoform (RNAPIIa) over the phosphorylated RNAPII isoform (RNAPIIo), therefore inhibiting transcription (Cheng & Price, 2007; Chodosh et al., 1989; Marshall & Price, 1992). Results in Figure 4A show that even when RNAPIIa greatly outnumbers RNAPIIo, a clear reduction in RPB1 levels is still observed upon UV irradiation, suggesting that transcriptionally inactive RPB1 molecules can be targeted for proteasomal degradation. Next, we used specific inhibitors for two of the main CTD kinases involved in transcriptional regulation, cyclin dependent kinases 7 and 9 (CDK7 and CDK9). CDK7, part of TFIIH, phosphorylates RPB1’s CTD residues at positions serine 5 and favors transcriptional initiation, while CDK9, part of the pTEFb complex, phosphorylates serines at position 2, a hallmark of productive elongation (Egloff et al., 2012). UV irradiation of WT HaCaT cells treated with CDK7’s inhibitor THZ1 or CDK9’s inhibitor BAY1251152 showed similar results to those obtained with DRB: a clear reduction in RPB1 levels in response to UV exposure (Figure S4A). To further confirm that RPB1’s phosphorylation state and transcriptional activity are not a pre-requisite for degradation, we took advantage of a mutant version of RPB1 that cannot be phosphorylated at serines 2 and 5 because of alanine replacement (A2A5, alanine 2 and 5 mutant polymerase) (M. J. Muñoz et al., 2009). Results in Figure 4B and S4B showed that A2A5 mutant RPB1 molecules are also degraded upon UV exposure, favoring the notion that transcriptionally inactive RPB1 molecules can be targeted for degradation. In this sense, the decrease in the unphosphorylated RNAPIIa isoform band in UV treated cells was thus far interpreted as a conversion into the phosphorylated RNAPIIo isoform, which would later be degraded. In view of the results shown here, the observed disappearance of the RNAPIIa band may simply reflect its degradation.

**Figure 4.**
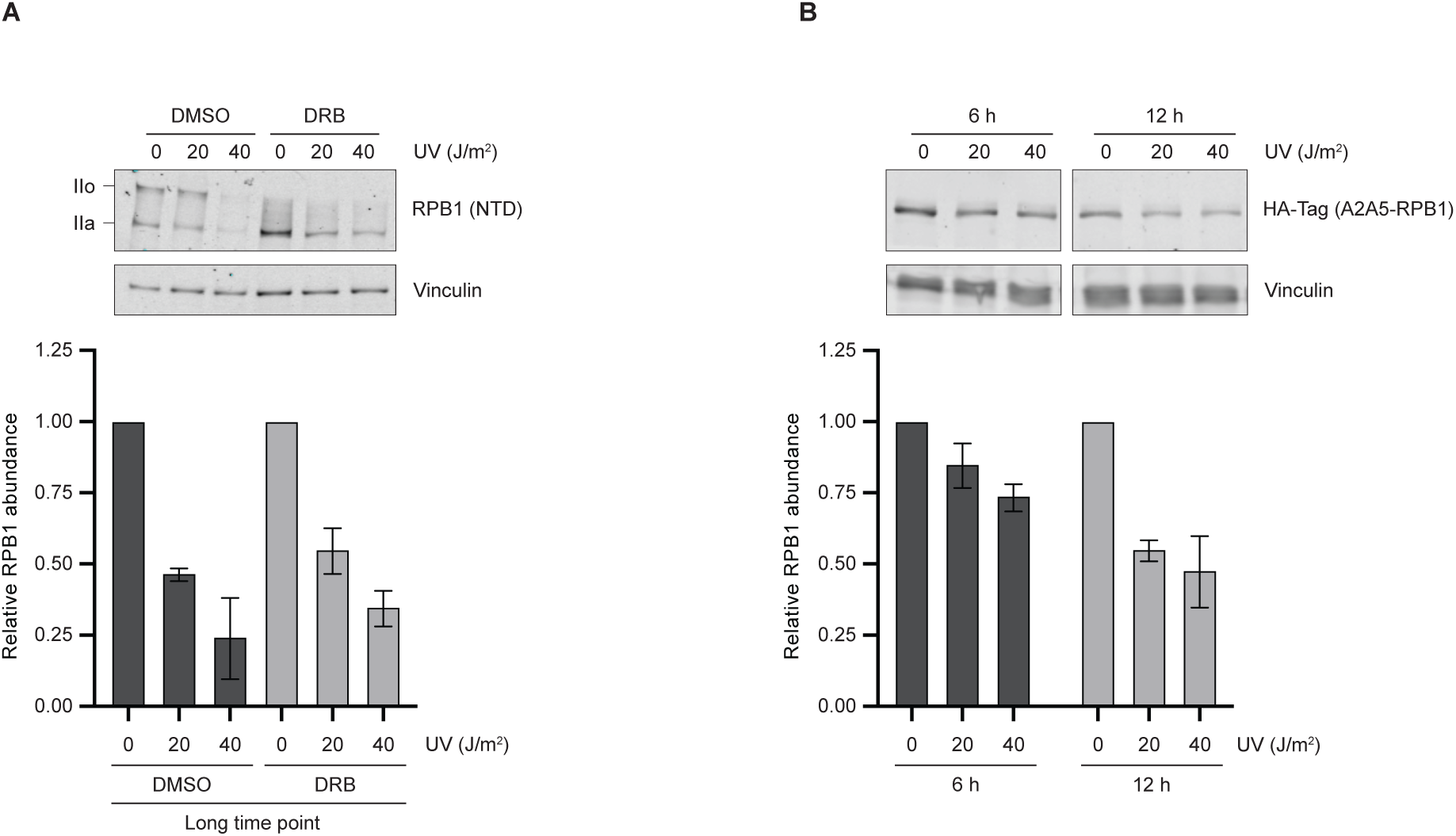
Transcriptionally inactive RPBl molecules can be targeted for degradation. (A) G1 arrested WT human keratinocytes were incubated for 3 h with the kinase inhibitor ORB (100 µM) and then irradiated with the indicated doses of UV light. Cells were harvested 10 h after irradiation. llo and lla indicates the phosphorylated and unphosphorylated forms of RPB1. Relative RPB1 abundance was assessed as before. lmages of a representative experiment and mean ± SEM of RPB1 abundance relative to 0 J/m^2^ condition from two independent experiments are shown. (B) HEK293T human cells were transfected with a plasmid encoding an HA-tagged version of RPB1 that cannot be phosphorylated at serines 2 and S because of alanine replacement (A2AS-RPB1). 48 h after transfection, cells were irradiated with the indicated doses of UV light and harvested after 6 h or 12 h. Relative levels of HA-tagged A2AS-RPB1 were assessed by western blot using antibodies against the HA tag and Vinculin as a loading control. lmages of a representative experiment and mean ± SEM of HA-tagged A2AS-RPB1 abundance relative to 0 J/m^2^ condition from two independent experiments are shown.

### NER-dependent degradation of RPB1 depends on Cullin-RING Ligases

Different E3 ubiquitin ligases were shown to be involved in RPB1 proteasomal degradation (J. C. Muñoz et al., 2022). Relevant for DNA repair, Cullin4 RING ubiquitin ligases (CRL4s) complexes were shown to be extensively remodeled in response to UV irradiation (Reichermeier et al., 2020), and CRL4^CSA^, an E3 complex associated with TC-NER, was shown to partially control RPB1 degradation (Tufegdžić Vidaković et al., 2020). Moreover, and in relation to GG-NER, our results suggest that CRL4^XPE/DDB2^ is also involved in the control of RPB1 degradation. Results in Figure 2 and S2A showed that dKO cells, as well as single XPE knockout cells, displayed reduced degradation levels of RPB1 in response to damage. Thus, to gather additional evidence supporting a role for CRLs in RPB1 degradation, we used the neddylation inhibitor MLN4924, which impedes NEDD8 conjugation to CRLs and their subsequent activation. Results in Figure 5A show that in HaCaT cells exposed to UV light, RPB1 degradation greatly depends on CRL activity. Moreover, TC-NER dependent RPB1 degradation, induced by Illudin S, was also CRL-dependent (Figure 5B) and the same was observed for degradation of transcriptionally inactive RPB1 molecules, as in the case of DRB and UV combined treatments (Figure 5C). These results suggest that CRL activities related to TC-NER and GG-NER are responsible for the control of RPB1 degradation in human keratinocytes.

**Figure 5.**
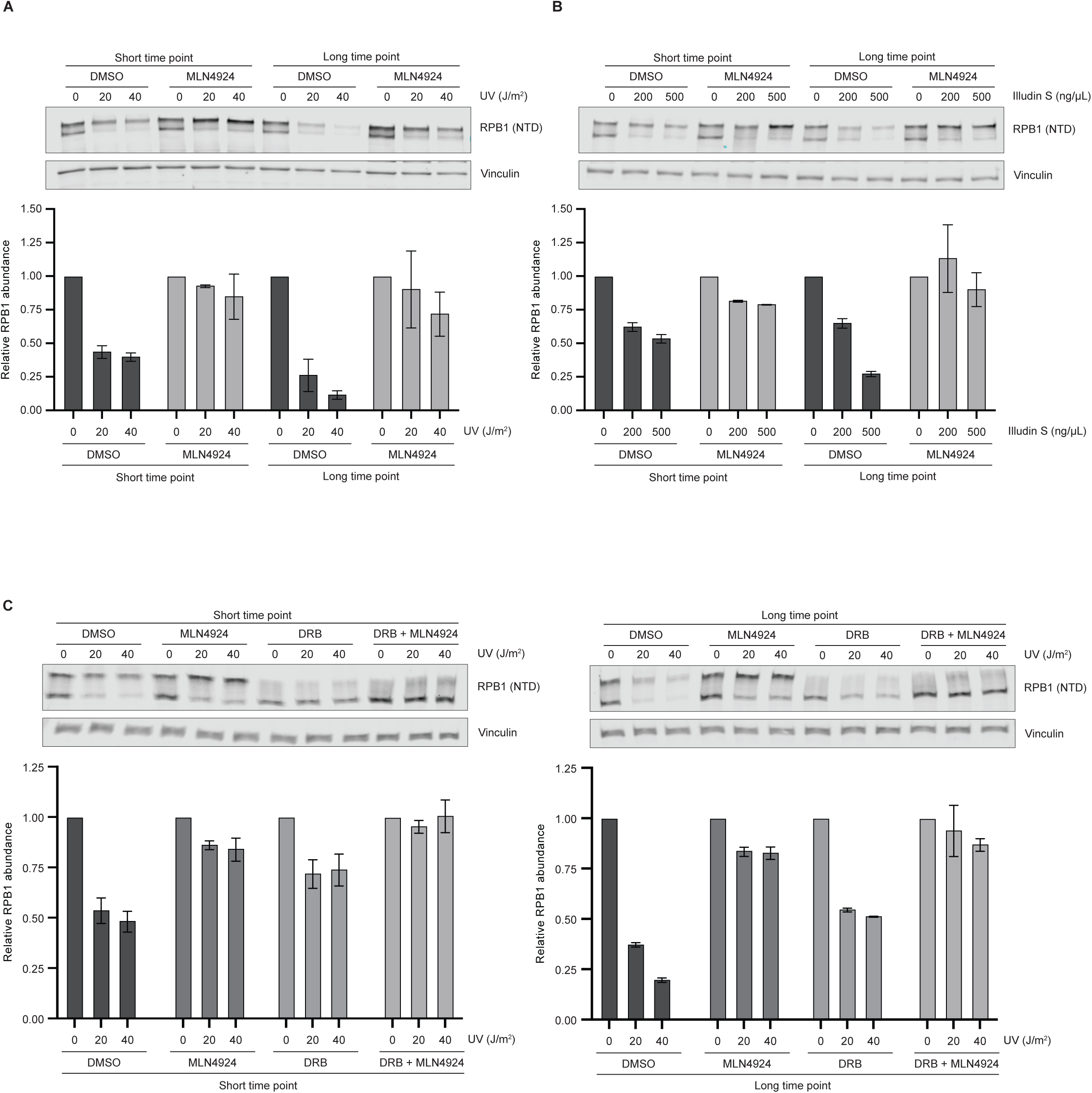
NER-related CRL4 E3 ubiquitin ligases target RPBl degradation. (A) Gl arrested WT human keratinocytes were incubated for l h with the neddylation inhibitor MLN4924 (l0 µM) before irradiation with the indicated doses of UV light. Cells were harvested 3 h (short time point) or l2 h (long time point) after irradiation and the relative RPBl abundance was assessed as before. Images of a representative experiment and mean ± SEM of RPBl abundance relative to 0 J/m^2^ condition from two independent experiments are shown. (B) Gl arrested WT human keratinocytes were incubated for l h with the neddylation inhibitor MLN4924 (l0 µM) before adding the indicated doses of Illudin S. Cells were harvested after 3 h (short time point) or l2 h (long time point). Relative RPBl abundance was assessed as before. Images of a representative experiment and mean ± SEM of RPBl abundance relative to 0 J/m^2^ condition from two independent experiments are shown. (C) Gl arrested WT human keratinocytes were incubated with the neddylation inhibitor MLN4924 for l h (l0 µM), with the kinase inhibitor DRB (l00 µM) for 3 h, or a combination of both, before irradiation with the indicated doses of UV light. Cells were harvested 3 h (short time point) or l2 h (long time point) after irradiation. Relative RPBl abundance was assessed as before. Images of a representative experiment and mean ± SEM of RPBl abundance relative to 0 J/m^2^ condition from two independent experiments are shown.

### Proposed model for RPB1 degradation in response to DNA damage

Based on the results presented here, we propose the model shown in Figure 6, illustrating a new role for NER in the control of RPB1 levels and, therefore, RNAPII activity. Lesion recognition by TC-NER (short time points, left panel) or GG-NER (longer time points, right panel) not only induces lesion repair but also CRL-dependent RPB1 ubiquitylation and degradation. In agreement with an *in trans* degradation pathway, transcriptionally inactive RBP1 molecules can be targeted for degradation, strongly suggesting that RPB1 degradation in response to damage is not related to lesion repair, but to the control of gene expression and cell fate under stress.

**Figure 6:**
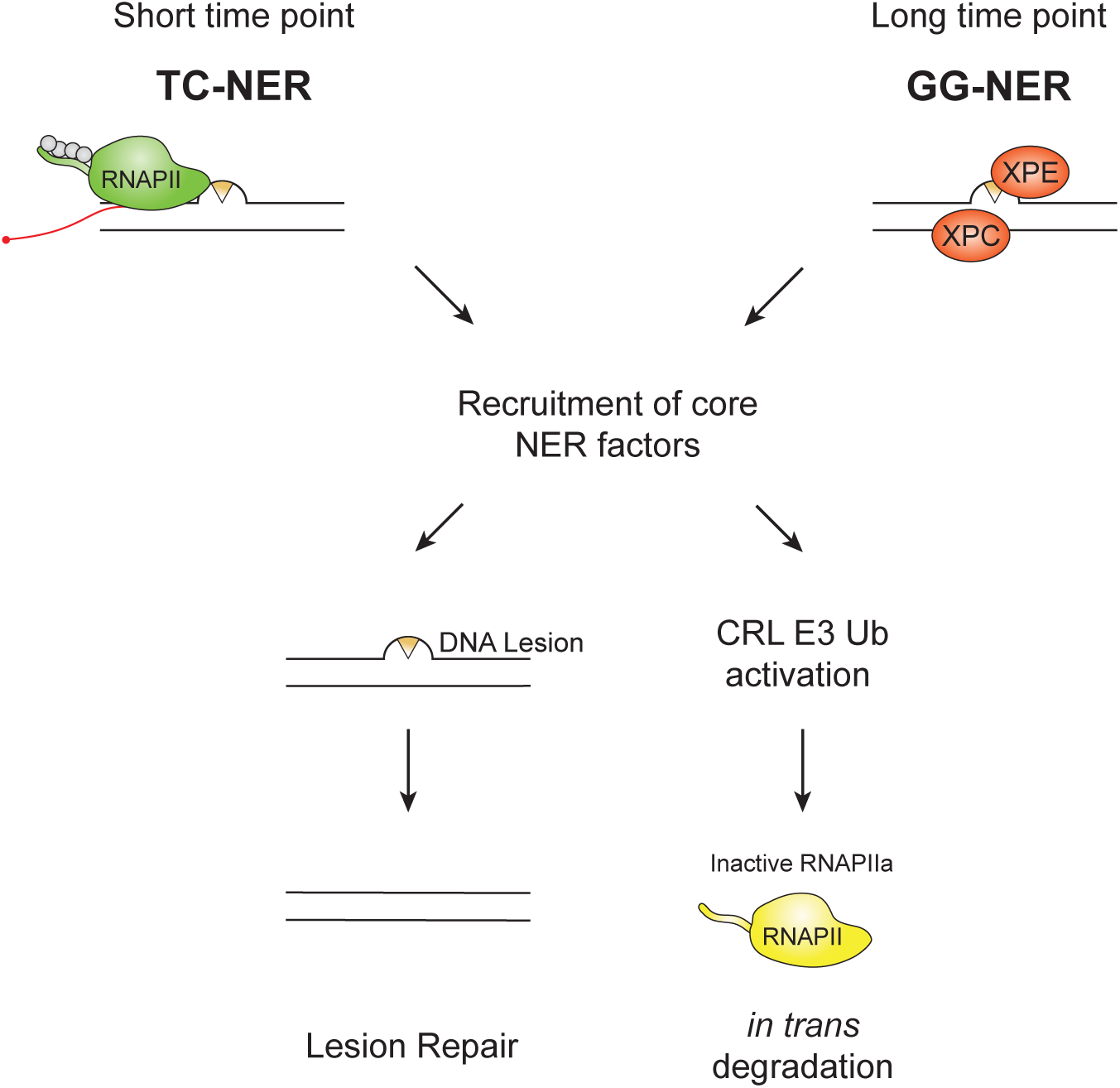
Model for the control of RPB1 levels by the NER system. The proposed model for the control of RNAPII activity under genotoxic stress highlights the role of NER not only in DNA repair, but also in the control of RPB1 degradation. Immediately after damage induction, lesion recognition will mainly depend on TC-NER (short time points). Thereafter, as transcription is inhibited, GG-NER will deal with most of damage recognition (long time points). Irrespectively of how the lesion was recognized, recruitment of core NER factors induces lesion repair as well as activation of E3 ubiquitin ligases able to target, *in trans*, transcriptionally inactive RPB1 molecules for ubiquitylation and proteasomal degradation.

## Discussion

### NER connects repair and gene expression

In this work we have identified a NER-dependent pathway that controls RPB1 levels and connects two major DNA-based events: lesion repair and gene expression. As can be seen throughout this work and elsewhere, more damage implies less RPB1 levels. Consequently, these findings provide a rationale for the control of gene expression under stress: the number of lesions determines the amount of repair that, through the control of RPB1 abundance, modulates the gene expression response.

RNAPII fate in response to damage is controlled sequentially, first by TC-NER and then by GG-NER, the latter being greatly responsible in the control of RPB1 levels in human keratinocytes exposed to UV light. Apart from the results showed here, the sequential control of RNAPII levels is supported by the fact that transcription is inhibited shortly after damage and, therefore, lesion recognition by TC-NER will decrease consequently. Conversely, lesion recognition by GG-NER is active for longer periods of time, favoring the proposed sequential control of RPB1 abundance.

A retrospective analysis of relevant published data suggests that the NER-dependent pathway controlling RPB1 levels herein proposed might have been overlooked. The seminal work by Ratner and co-workers (Ratner et al., 1998) presented different clues suggesting that NER deficient cells have altered RPB1 levels in response to UV light: i) Results in figure 1 show that XPD mutant fibroblasts have reduced levels of RPB1 in comparison to WT cells, ii) Results in figure 6 show the same, but this time using XPA and XPG mutant fibroblasts and, iii) Figure 6 also shows that XPC mutant fibroblasts present higher levels of RPB1 than its WT counterpart (please see (Ratner et al., 1998)). Additionally, the Sancar laboratory showed an enhancement in template-strand repair in XPC mutant fibroblasts, a result that suits the idea of the present work, where a GG-NER deficient context leads to more RPB1, hence favoring transcription and template-strand repair (Hu et al., 2015). Therefore, we propose there is an enhancement of TC-NER activity in GG-NER deficient cells, not only because there is no repair by GG-NER, but also because of higher levels of RPB1 in response to any damage dealt by NER. Conversely, mutant cells in core NER repair factors (XPA, XPB, XPD, XPF and XPG) will deal with reduced levels of RPB1 and lower TC-NER activity. In line with this, different reports showed altered gene expression patterns in NER deficient cells in response to damaging agents (Andrade-Lima et al., 2015; M. J. Muñoz et al., 2017).

### *In situ* vs *in trans* degradation of RPB1

There is a substantial difference between *in situ* degradation or an *in trans* signaling cascade controlling RPB1 levels: i*n situ* degradation facilitates lesion clearance and repair while, in a signaling cascade, RPB1 degradation is not related to lesion clearance nor repair, but to the overall control of the gene expression response. In the latter scenario, lesion clearance and repair, as well as RPB1 degradation, would depend on RPB1 eviction from chromatin. In support of this, the ATP dependent segregase VCP/p97, able to regulate the dynamic interaction of RPB1, XPC and XPE/DDB2 with chromatin, showed to be necessary for the control of RPB1 levels in response to damage (Ribeiro-Silva et al., 2020; Steurer et al., 2022). Moreover, RNAPII was also shown to be released from the DNA template during TC-NER (Chiou et al., 2018). On the contrary, *in situ* degradation of lesion stalled RPB1 molecules, as suggested by the last resort model, implies the direct interaction between the proteasome and the chromatin-engaged transcriptional complex. In this sense, interaction between the 26S proteasome and DNA has been reported in yeast (Auld et al., 2006; Gillette et al., 2004) and in humans (Epanchintsev et al., 2017; Kito et al., 2020), supporting a model in which clearance of the lesion can be achieved directly by the proteasome. It is then possible that *in situ* and *in trans* degradation co-exist. UV light, one of the most common cytotoxic agents used in research, induces different types of lesions that might well condition the pathway of choice, i.e., *in situ* or *in trans* degradation. Apart from pyrimidine dimers and many others, UV light exposure favors the crosslink between aromatic amino acids and nucleic acids (Greenberg, 1979). In this sense, cryo-electron microscopy data of an elongating RNAPII complex showed that two phenylalanine and six tyrosine residues are sufficiently close, less that ten angstroms, to a DNA or RNA nucleotide (protein data bank accession number 8B3D). We hypothesize that a crosslink between RPB1’s residues and DNA or RNA may represent a scenario in which *in situ* degradation is favored over chromatin eviction.

### Transcriptionally inactive RPB1 molecules are degraded

Regarding the nature of the RPB1 molecule that is degraded by the proteasome, i.e., phosphorylated or not, the conceptual framework thus far pointed towards RPB1 molecules that were part of active RNAPII complexes. Nevertheless, results in Figure 4 clearly show that transcriptionally inactive RPB1 molecules are degraded. In this sense, degradation of promoter-proximal paused RPB1 molecules (Bay et al., 2022; Steurer et al., 2022) might well be interpreted as degradation of transcriptionally inactive or paused molecules, although in this case enriched in phosphorylated serine 5. Together with results in Figure 4 showing that unphosphorylated RPB1 molecules are still degraded, it is then conceivable that RPB1 molecules effectively targeted for degradation are transcriptionally inactive with none or few nucleotides of transcribed RNA, as it would be the case for promoter proximal paused RPB1 molecules.

Recently, Steurer and co-workers (Steurer et al., 2022) suggested that degradation of promoter-proximal paused RPB1 is GSK3 dependent and DNA repair independent, opposite to what we have described throughout this work. Our results suggest a NER dependent degradation pathway of unphosphorylated RPB1 molecules that is also active in the presence of GSK3 inhibitors. Given the complexity of human cells and the relevance of the factor of study (RPB1 levels / RNAPII activity), it is expected that different mechanisms are involved in the control of RPB1 levels. As commented before, the *POLR2A* gene ranked at the top of the genes whose copy-number is of paramount importance for proper cell function. Therefore, it is not surprising that different mechanisms, active in different cell types or states, help to determine the abundance of RPB1. For the case herein described, the fact that NER modulates RBP1 levels suggest that RPB1 abundance might be controlled not only under an acute response to damage, but also in normal conditions. NER deals with different types of lesions, including some derived from oxidative metabolism like 5′,8-cyclopurines (Kropachev et al., 2014), and therefore, it is possible that high-energy demand tissues of NER-deficient patients present abnormal levels of RPB1 abundancy and, in turn, misregulation of gene expression.

### Transient activation of NER-related Cullin Ring-Ligases

In relation to the E3 ubiquitin ligases involved, our results suggest neddylation dependent CRL E3 ubiquitin ligases associated with TC-NER or GG-NER, i.e., CRL4^CSA^ and CRL4^XPE/DDB2^, are involved in the control of RPB1 stability. The fact that defects in the completion of the repair reaction are associated with an enhancement in RPB1 degradation (Figure 3), suggests that CRL E3 ligases are transiently activated during the NER reaction, and that such an activation continues until DNA structure is restored, only after DNA synthesis and ligation. Supporting a model of a transient activation of E3 complexes after damage, it has been recently published that CRL4 complexes are dynamically reorganized upon UV exposure (Reichermeier et al., 2020).

### RPB1 as a limiting factor under stress

The idea of an intimate connection between DNA repair and gene expression was established many years ago, when it became apparent that certain factors play roles in both processes. Thus, the notion of a dual and limiting factor operating in repair but not in transcription was conceived in an effort to comprehend the global inhibition of gene expression following DNA damage. Based on the findings presented in this study, we propose another layer in the coupling between gene expression and DNA repair, in which NER-mediated repair turns RPB1 into a limiting factor, restricting RNAPII activity and, hence, shaping the gene expression response to DNA damage.

## Acknowledgments

We thank all the members of the MJM and MF groups for their patience and contribution to this work. The MJM group thanks Dr Ignacio Schor and Dr Manuel de la Mata for assistance with data sets analysis in figure S1A, Mr. Mariano Lopez Gringauz for technical assistance, M.S. Emilia Haberfeld for statistical analysis and Dr Luciana Giono for model drawing.

M.J.M laboratory is supported by grants from the Agencia Nacional de Promoción Científica y Tecnológica (ANPCyT, Argentina) to M.J.M (2020-1025) and to L.A.B. (2020-3194).

M.F. laboratory is supported by grants from Associazione Italiana per la Ricerca sul Cancro, AIRC, Italy, (AIRC-IG-21416) and Ministero dell’Istruzione, dell’Università e della Ricerca (MIUR-PRIN-2015SJLMB9).

## Author Contribution

Experiments were designed and performed by R.C, J.C.M, I.B, G.B, L.A.B and M.J.M. Conceptualization and writing were done by M.J.M. All the authors read and corrected the manuscript.

## Methods

### Cell culture

HaCaT human keratinocytes were cultured in 5% CO_2_ incubators at 37°C in DMEM (ThermoFisher Scientific #10566016) supplemented with 10 % FBS (Thermo Fisher Scientific #10270098), and 1% Primocin (Invivogen #ant-pm-1). HEK293T were cultured in 5% CO_2_ incubators at 37°C in DMEM supplemented with 10% FBS (Thermo Fisher Scientific #10270098) and 1% Pen-Strep (GIBCO #XXXXX). HaCaT cells were routinely sub-cultured two to three times a week using a 1:4 dilution ratio while HEK293T were sub-cultured using a 1:10 dilution ratio. All cell lines were proved to be mycoplasma-free.

### Drug treatments

When indicated, cells were treated with the following drugs: Illudin S (MedChemExpress #HY-125098); MG132 (Selleckchem #S2619); Spironolactone (Selleckchem #S4054); α-amanitin (Sigma #A2263); Aphidicolin (MedChemExpress #HY-N6733); ATR inhibitor VE822 (Selleckchem # S7102); ATM inhibitor KU60019 (Selleckchem # S1570); DNA-PK inhibitor NU7441 (Selleckchem #S2638); GSK3 inhibitor CHIR-99021 (Sigma); 5,6-Dichlorobenzimidazole 1-β-D-ribofuranoside (DRB) (SIGMA #D1916); CDK7 inhibitor THZ1 (Selleckchem #S7449); CDK9 inhibitor BAY1251152 (Selleckchem #S8730); MLN4924 (Selleckchem #S7109)

### Plasmid construction

The oligonucleotide duplex (AGCTTGCCGCCACCATGACTAGTTATCCTTACGATGTGCCAGACTACGCTAGCG; GATCCGCTAGCGTAGTCTGGCACATCGTAAGGATAACTAGTCATGGTGGCGGCA) containing the Kozak sequence and the HA epitope was inserted into HindIII/BamHI treated pcDNA5/FRT/TO vector (Thermo Fisher Scientific). To obtain HA tagged RPB1-A2A5 expression vector, the 1579 bp PvuI/NheI fragment spanning a section of the ampicillin selectable marker, CMV promoter and HA epitope coding sequence of this resulting plasmid was transferred to pEVRF1-RPB1-A2A5 (M. J. Muñoz et al., 2009) employing the same restriction enzymes. An equivalent vector for the expression of WT HA tagged RPB1 was derived replacing the A2A5 coding region of the former with the WT CTD equivalent contained in the XhoI/XbaI 2509 bp section of pEVRF1-RPB1 (M. J. Muñoz et al., 2009).

### Generation of stable cell lines

CRISPR-Cas9 single XPE KO clones of HaCaT cells were obtained as described in (M. J. Muñoz et al., 2017). CRISPR-Cas9 single XPC KO clones were obtained following the protocol described in (Ran et al., 2013). The gRNA for Cas9 targets within XPC first exon were cloned into the pSpCas9(BB)-2A-GFP (PX458), and stable transfectants were selected by Fluorescence-Activated Cell Sorting (FACS). The double KO XPC-XPE pools were obtained by doing the XPC KO in the XPE KO genetic background, and stable transfectants were selected by FACS followed by Puromycin selection. Single clones were obtained by clonal-density dilution. CRISPR-Cas9 KO clones for XPA were obtained in the same manner as XPC KO clones. Sequences for each specific gRNA are presented in Supplementary Table 1.

### G1-arrest

HaCaT human keratinocytes were seeded at 90% confluence in DMEM supplemented with 10% FBS and 1% Primocin. 24 h later, cells were washed twice with PBS 1X and the culture medium was changed to DMEM supplemented with 0.5% FBS and 1% Primocin. Cells were further incubated for 48 h to allow for G1-arrest.

### UV-irradiation

HaCaT and HEK293T cells were washed once with PBS 1X before irradiation with UVC light (254 nm). UV irradiation was performed with a CL-1000 Shortwave Crosslinker (UVP) or a custom-made UV-irradiation apparatus. UV light doses were quantified with an external UVC radiometer.

### siRNA Knockdown experiments

HaCaT human keratinocytes at 70% confluence (FBS 10%, Primocin 1%) were transfected using RNAiMAX (Thermo Fisher Scientific #13778075) according to manufacturer’s instructions. Pre-designed human siGenome pool siRNAs were obtained from Dharmacon. 24 h after transfection, when cells were at 100% confluence, culture medium was changed to DMEM supplemented with 0.5% FBS and 1% Primocin, after which cells were further incubated for 48 h. After this time, cell population was G1-arrested and siRNA-transfected 72 h before.

### Western blot

Protein samples were obtained by harvesting cells with Laemmli Buffer 2X (Tris-HCl pH 6.8 125 mM, β-Me 10%, SDS 4%, Glycerol 20%). Samples were subjected to SDS-polyacrylamide gel electrophoresis and transferred to a nitrocellulose membrane (GE Healthcare Life Sciences #10600002). Membranes were blocked at room temperature for 1 h with 5% milk in T-TBS (1% Tween 20 - Tris Buffered Saline buffer) and incubated overnight at 4 °C with the indicated primary antibodies diluted in 5% milk in T-TBS. Membranes were washed three times with T-TBS and incubated for 1h at room temperature with secondary antibodies. Membranes were washed again three times with T-TBS and visualized by using an Odyssey Imaging System for fluorescent secondary antibodies or incubated with a chemiluminescent substrate ECL reagent (Thermo Scientific Super Signal #34094) for HRP-conjugated secondary antibodies. For a list of antibodies used throughout this work please see Supplementary Table 1.

### 4sU Dot blot

4sU Dot Blot was performed following the protocol described in (Gregersen et al., 2020). Briefly, WT and dKO HaCaT cells were treated for 15 min with 4sU 1 mM prior to harvesting, and total RNA was extracted using TriPure reagent (Roche #11667157001) according to manufacturer’s instructions. Total RNA was quantified with an RNA-HS kit (Thermo Fisher Scientific #Q10210) and 5 µg of RNA per sample were biotinylated with MTSEA biotin-XX linker (Biotium #BT90066) for 30 min at room temperature in dark conditions. Biotinylated RNA was re-purified with Phenol:Chloroform:Isoamylic (25:24:1) (ThermoFisher #15593031) and immobilized onto a Nylon membrane (CATNUM) using a Dot Blot apparatus (Biorad #1706545). RNA was crosslinked to the membrane by irradiating with one pulse of 2000 J/m^2^ of UV Light. Membranes were incubated with blocking solution (SDS 10%, EDTA 500 mM, PBS1X) for 20 min at room temperature followed by incubation with a 1:50000 dilution of Alexa Fluor 790-Streptadivin (Jackson ImmunoResearch) #016-650-084). Membranes were washed multiple times with 1:10 serial dilutions of washing buffer (SDS 10%). Alexa Fluor 790-Streptadivin signal was acquired with an Odyssey Imager System. Lastly, membranes were washed once with H_2_Od and stained for 10 minutes with methylene blue solution (0.5% methylene blue, 0.5M sodium acetate) for loading control. De-staining was performed by washing the membranes with H_2_O overnight.

### RT-qPCR

Total RNA was extracted using TriPure reagent (Roche #11667157001) according to manufacturer’s instructions. RNA samples were treated with DNAse (Promega #M6101) and reverse transcription of DNase-treated RNA was performed with random decamers using SuperScript III reverse transcriptase (ThermoFisher Scientific #18080093). cDNA was subjected to qPCR using specific primers for each gene with MasterMix (ThermoFisher Scientific #4472908) following manufacturer’s instructions. qPCR cycles consisted of 40 cycles of 15 s denaturation at 95°C, 15 s annealing at 60 °C and 45 s for primer extension at 72 °C. Sequences for the primers used are shown in Supplementary Table 1.

### Southern blot

WT and dKO HaCaT cells were irradiated and harvested at the indicated time points for genomic DNA extraction with a QIAGEN QIAmp DNA Mini Kit (Qiagen #51304), according to manufacturer’s instructions. DNA samples were loaded in a 0.8% agarose gel and runed for 1 h at 100 V. The agarose gel was then transferred by southern blot to a Hybond+ membrane (Amersham Hybond-XL GE Healthcare) with SSC10X buffer (1.5 M NaCl, 0.15 M Tris-Na Citrate, pH 5) and DNA was crosslinked by baking the membrane for 2 h at 80 °C. The membrane was then blocked for 1 h at room temperature with 5% milk in T-PBS (1% Tween 20 – PBS1X), washed once with T-PBS and incubated with primary antibodies at 4 °C overnight. For a list of antibodies used throughout this work please see Supplementary Table 1. Primary antibodies were prepared in BSA 5% T-PBS. Membranes were then washed three times with TENT buffer (20 mM Tris HCl pH 8 20 mM, 137 mM NaCl, Tween 0,1%) and incubated for 1 h at room temperature with secondary antibodies. Membranes were washed again three times with TENT buffer and incubated with a chemiluminescent substrate ECL reagent for HRP-conjugated detection of secondary antibodies.

### FACS analysis

WT and dKO HaCaT cells were harvested by gently scrapping the plates in PBS 1X and centrifuged at 300 g for 5 minutes. PBS1X was aspirated and cells were resuspended in ethanol 70% by up and down pipetting to avoid cell aggregation. Cells were then incubated for 2 h at 4°C for fixation followed by a centrifugation step of 5 min at 300 g. The remaining ethanol was washed once with PBS1X and cell pellets were resuspended in 1 mL of propidium iodide solution (propidium iodide 120 mM, 0.1% Triton X-100, 100 ng/µL RNaseA, PBS1X) before incubation for 30 minutes at room temperature. Samples were then vortexed and cell fluorescence was measured with a FACS Aria II flow cytometer. Data were analyzed using Flow Jo software.

---

**Supplementary Figure 1, related to Figure 1.**
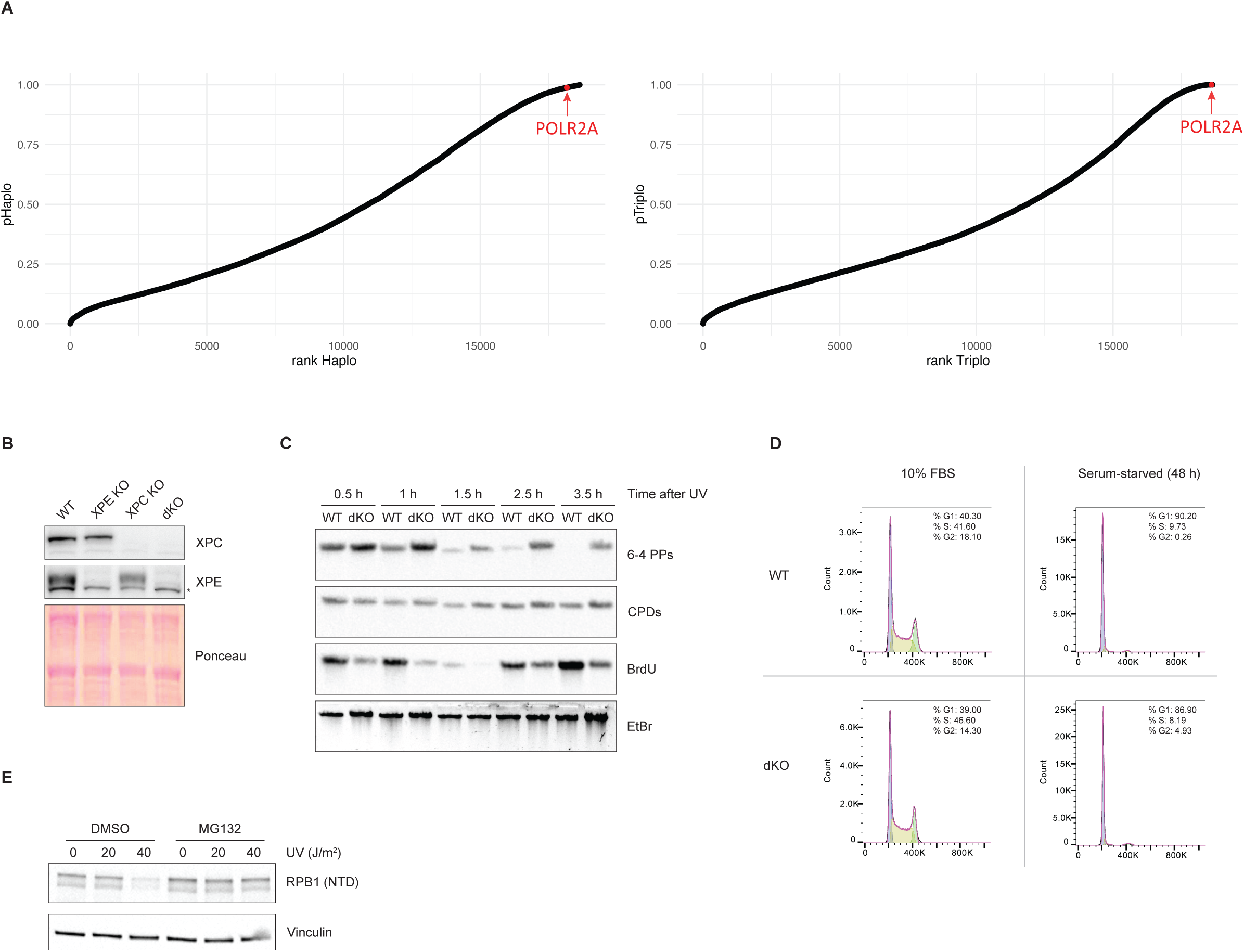
TC-NER controls RPB1 levels in human keratinocytes. (A) Predicted dosage sensitivity at a single gene resolution. The Y axis represents the probability of developing a disease when any given human gene (ranked in the X axis) is haploid (haploinsufficiency, left panel) or has one extra copy (triplosensitivity, right panel). POLR2A, the gene encoding RPB1, is shown in red. Data sets were obtained from (Collins et al., 2022). (B) Western blot showing the levels of XPC and XPE in WT, XPE KO, XPC KO and XPC-XPE double KO (dKO) HaCaT human keratinocytes. Ponceau staining is shown as a loading control. The asterisk in XPE western blot indicates an unspecific band. (C) G1 arrested WT and XPC-XPE double KO (dKO) HaCaT cells were irradiated with 10 J/m^2^ of UV light and harvested at the indicated time points. 1 h before harvesting, BrdU (10 µM) was added to culture medium. Genomic DNA was then purified and subjected to southern blot analysis. Membranes were probed with antibodies against Cyclobutane Pyrimidine Dimers (CPDs), 6-4 Pyrimidine-pyrimidone Photoproducts (6-4PPs) and BrdU. Ethidium Bromide (EtBr) staining of total DNA is shown as loading control. (D) WT and XPC-XPE double KO (dKO) HaCaT human keratinocytes were incubated in normal conditions (10% FBS) or grown until 100% confluence before being serum-starved for 48 h to induce G1 arrest (cell cycle arrest protocol, see Methods). Samples were harvested, fixed and stained with propidium iodide (PI) for flow cytometry analysis of DNA content. Cell cycle histograms and the percentage of cells on each cell cycle stage are shown. The experiment was performed twice, and a representative experiment is shown. (E) G1 arrested WT HaCaT human keratinocytes were incubated with the proteasome inhibitor MG132 (10 µM) for 1 h before irradiation with the indicated doses of UV light. Cells were harvested 10 h after. Relative RPB1 abundance was assessed by western blot using antibodies against the N-Terminal Domain (NTD) of RPB1 and Vinculin as a loading control. A representative experiment of at least two independent experiments is shown.

**Supplementary Figure 2, related to Figure 2.**
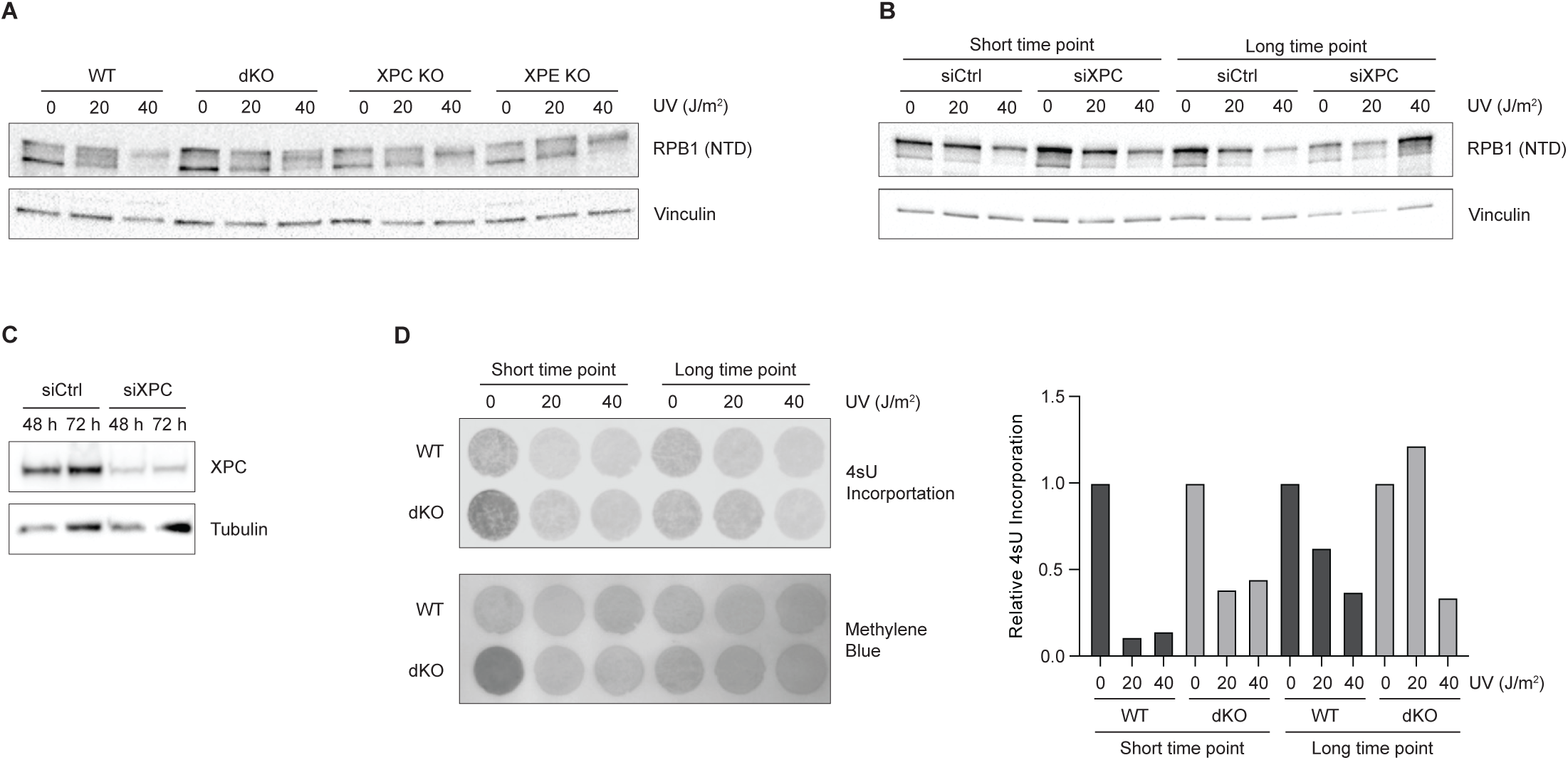
GG-NER controls RPB1 levels in human keratinocytes. (A) G1 arrested WT, XPC-XPE double KO (dKO clone #2), XPC KO and XPE KO HaCaT cells were irradiated with the indicated doses of UV light and harvested 12 h after (long time point). Relative RPB1 abundance was assessed as before. A representative experiment of at least two independent experiments is shown. (B) WT HaCaT human keratinocytes were transfected with scramble siRNAs (siCtrl) or siRNAs directed against XPC (siXPC). 24 h later, cells were arrested in G1 by serum withdrawal for 48 h and then irradiated with the indicated doses of UV light. Cells were finally harvested 3 h (short time point) or 12 h (long time point) after irradiation. Relative RPB1 abundance was assessed as before. A representative experiment of at least two independent experiments is shown. (C) WT HaCaT human keratinocytes were transfected with siCtrl or siXPC siRNAs and harvested 48 h or 72 h after. siRNA-mediated knock down of XPC was assessed by western blot using antibodies against XPC and Tubulin as a loading control. Images of a representative experiment are shown. (D) G1 arrested WT and XPC-XPE double KO (dKO) HaCaT cells were irradiated with the indicated doses of UV light and harvested 3 h (short time point) or 12 h (long time point) after. 15 minutes before harvesting, cells were treated with a pulse of 4sU (1 mM) and total RNA was then prepared. Samples were subjected to 4sU dot blotting, showing global nascent RNA. Methylene blue staining of total RNA is shown as loading control. Left panel: 4sU and Methylene Blue membranes. Right panel: 4sU incorporation levels relative to total RNA.

**Supplementary Figure 3, related to Figure 3.**
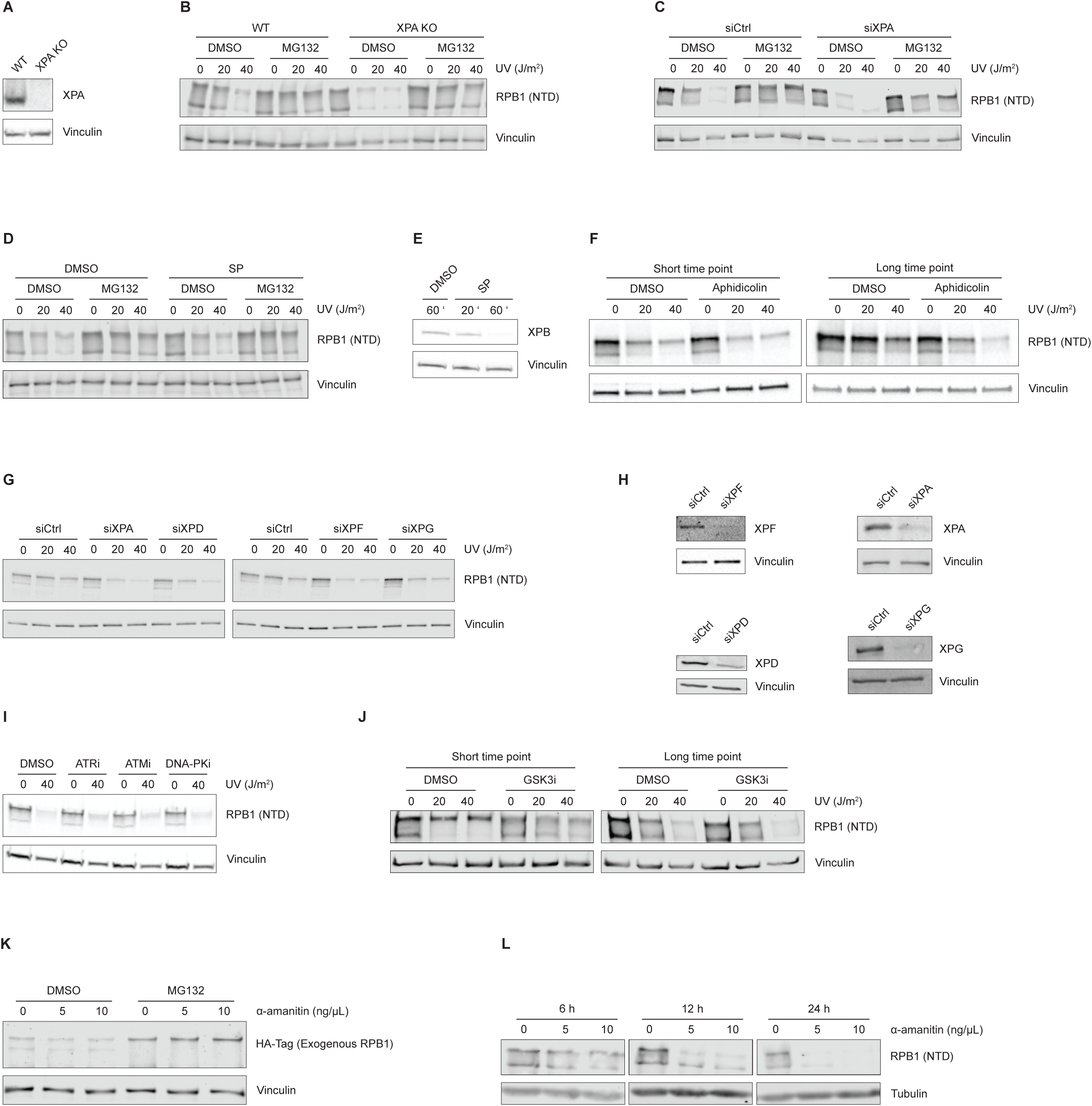
NER-mediated DNA repair controls RPB1 degradation in trans. (A) Western blot showing XPA relative levels in WT and XPA KO HaCaT cells. Vinculin was probed as loading control. (B) G1 arrested WT and XPA KO HaCaT cells were incubated for 1 h with the proteasome inhibitor MG132 (10 µM), then irradiated with the indicated doses of UV light and finally harvested 12 h after. Relative RPB1 abundance was assessed as before. A representative experiment of at least two independent experiments is shown. (C) WT HaCaT cells were transfected with scrambled (siCtrl) or XPA-specific (siXPA) siRNAs and further incubated for at least 72 h to allow mRNA knockdown and G1 arrest. Cells were then treated for 1 h with the proteasome inhibitor MG132 (10 µM), irradiated with the indicated doses of UV and finally harvested 12 h after. Relative RPB1 abundance was assessed as before. A representative experiment of at least two independent experiments is shown. (D) To induce XPB knockdown, G1 arrested WT HaCaT cells were incubated for 1 h with Spironolactone (2.5 µM). When indicated, the proteasome inhibitor MG132 was also added (10 µM). Cells were then irradiated with the indicated doses of UV light and harvested after 10 h. Relative RPB1 abundance was assessed as before. A representative experiment of at least two independent experiments is shown. (E) G1 arrested WT HaCaT human keratinocytes were incubated with Spironolactone (SP, 2.5 µM) for 20 or 60 minutes and then harvested for western blotting. XPB abundance was assessed using antibodies against XPB and Vinculin as loading control. A representative experiment of at least two independent experiment is shown. (F) G1 arrested WT HaCaT human keratinocytes were incubated for 2 h with the DNA polymerase inhibitor Aphidicolin (300 nM) and then irradiated with the indicated doses of UV light. Cells were harvested after 3 h (short time point) or 12 h (long time point). Relative RPB1 abundance was assessed as before. A representative experiment of at least two independent experiments is shown. (G) WT HaCaT human keratinocytes were transfected with scrambled (siCtrl) or specific (siXPA, siXPD, siXPF and siXPG) siRNAs and further incubated for 72 h to allow for mRNA knockdown and G1 arrest. Cells were then irradiated with the indicated doses of UV light and harvested 12 h later. Relative RPB1 abundance was assessed as before. A representative experiment of at least two independent experiments is shown. (H) WT HaCaT human keratinocytes were transfected with siRNAs directed against XPA, XPD, XPF, and XPG and harvested after 72 h. Knockdown of each XP factor was assessed by western blot using specific antibodies and Vinculin as loading control. Representative images are shown. (I) G1 arrested WT HaCaT human keratinocytes were incubated with ATR (VE822 1 µM), ATM (KU60019 3 µM), or DNA-PK (NU7441 1 µM) inhibitors for 1 h, irradiated with 40 J/m^2^ and harvested 12 h after. The relative abundance of RPB1 was assessed as before. A representative experiment of at least two independent experiments is shown. (J) G1 arrested WT HaCaT human keratinocytes were exposed to the indicated doses of UV light. Immediately after irradiation, the GSK3 inhibitor CHIR-99021 (20 µM) was added, and cells were harvested 3 h (short time point) or 12 h (long time point) after. Relative RPB1 abundance was assessed as before. A representative experiment of at least two independent experiments is shown. (K) HEK293T cells were transfected with a plasmid encoding an HA-tagged, α-amanitin resistant version of RPB1. 48 h after transfection, cells were incubated for 1 h with the proteasome inhibitor MG132 (10 µm) and then with the indicated doses of α-amanitin. Cells were harvested 12 h after. Relative levels of the exogenous HA-tagged and α-amanitin resistant version of RPB1 were assessed by western bot using antibodies against the HA tag and Vinculin as a loading control. A representative experiment of at least two independent experiment is shown. (L) HEK293T human cells were treated with the indicated doses of α-amanitin and harvested after 6 h, 12 h or 24 h. Relative RPB1 abundance was assessed as before. A representative experiment of at least two independent experiments is shown.

**Supplementary Figure 4, related to Figure 4.**
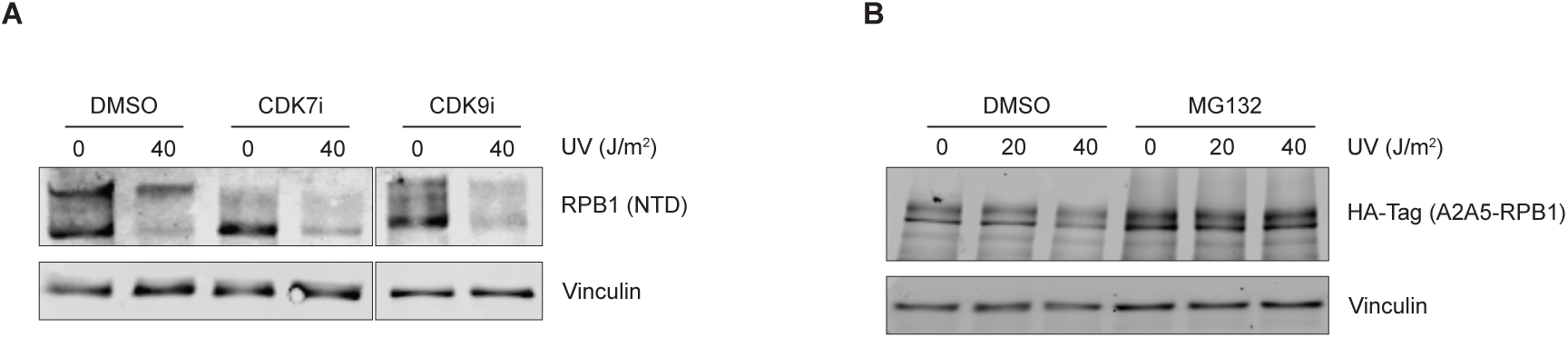
Transcriptionally inactive RPB1 molecules can be targeted for degradation. (A) G1 arrested WT human keratinocytes were irradiated with 40 J/m^2^ of UV light. Immediately after irradiation, the CDK7 inhibitor THZ1 (100 nM) or the CDK9 inhibitor BAY 1251152 (400 nM) was added, and cells were harvested 12 h after. Relative RPB1 abundance was assessed as before. A representative experiment of at least two independent experiment is shown. (B) HEK293T human cells were transfected with a plasmid encoding an HA-tagged version of RPB1 that cannot be phosphorylated at serines 2 and 5 (A2A5-RPB1). 48 h after transfection, cells were incubated with MG132 (10 µM) for 1 h before irradiation with the indicated doses of UV light and finally harvested 12 h after. Relative levels of the HA-tagged A2A5-RPB1 were assessed by western blot using antibodies against the HA tag and Vinculin as a loading control. The experiment was performed twice, and images of a representative experiment are shown.

**Supplementary Table 1.**
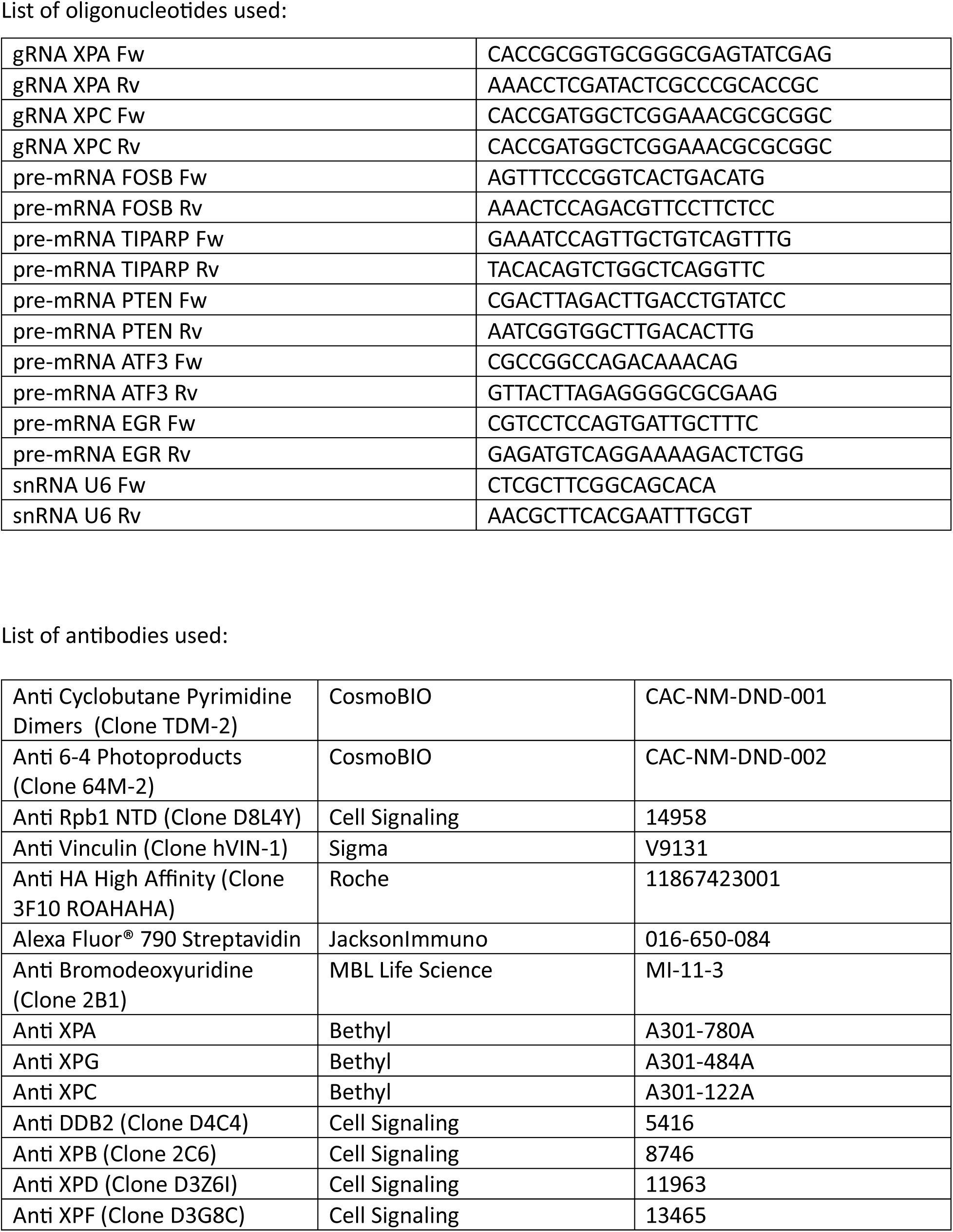

